# Niche construction: the case of industrial vinegar production

**DOI:** 10.1101/2025.01.02.631129

**Authors:** Reginald D. Smith

## Abstract

Niche construction is the process by which changes made to an environment by organisms also in turn drive their own evolution. While many examples of niche construction in the wild or human societies have been described, it is rare that a clear example of niche construction in an environment under human control can be demonstrated. Industrial vinegar production is a fermentation conducted by acetic acid bacteria. Importantly, the acetic acid bacteria produce acetic acid whose increasing acidity selects out acetic acid bacteria with lower tolerances for acidity. The remaining acetic acid bacteria continue to produce higher levels of acidity causing a directional selection that drives the acidity tolerance of the overall population higher while reducing the population’s genetic diversity. This niche construction will be described and modeled and its implications for the probable self-limiting constraints of niche construction in wild populations is discussed.

## 1 Introduction

Niche construction is a concept whereby the relationship between organisms and environment is modeled as a feedback between an organism which modifies its environment in a way that exerts selection pressure and influences the evolution of the same over time. The term was first coined by Odling-Smee [1–3]. Though its scientific and philosophical antecedents stretch to the earliest days of evolutionary biology and even further, the development niche construction was partly in response to a proposal of a “constructive” paradigm of organism and environment interaction proposed by Lewontin [4]. In this work, Lewontin proposed that the concept of the influence of environment on an organism depart from an open loop interpretation where the environment is a given that drives the organism’s adaptation, to a closed loop interpretation where the organism adapts to the environment but also alters it. The general equations to outline this notion are below in the continuous time differential equation as well as discrete time difference equation forms. Below *O* stands for the organism genotype and *E* for the environment. The phenotype is a interplay of *O* and *E*.

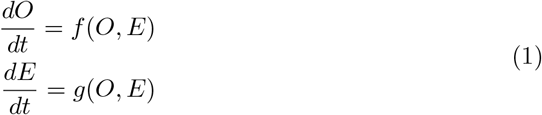

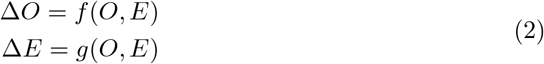

One key aspect of niche construction is the necessity of at least two loci to explain its effects [2]. One locus is responsible for the organism phenotype that alters the environment while a second locus evolves in response to this changed environment. Therefore the selective pressure on the second locus is indirectly driven by the first though there is no formal interaction such as epistasis.

While most studies of niche construction deal with macroscopic organisms whose behavior drives the alteration of the environment, niche construction is a valid concept for microscopic organisms as well, even the simple prokaryotes. Recent papers, have described several mechanisms by which bacteria engage in niche construction including *E. Coli* on a single carbon source growth medium producing secondary metabolites that are used as a nutrient source by other *Escherichia coli* driving population diversity [6, 7], human-bacteria feedback where antibiotic use by humans and the response by bacteria drive niche construction [8], niche construction for acidity tolerance by the ulcer causing bacteria *Helicobacter pylori* [9], and a general review of the types of niche construction in bacteria [10].

In this paper we will propose and model niche construction amongst acetic acid bacteria, the organisms that are responsible for the fermentation of vinegar. Separate genes are responsible for the production of acetic acid from ethyl alcohol (alcohol dehydrogenase and acetaldehyde dehydrogenase) and acidity tolerance (citrate synthase among others). The increase in acidity in vinegar fermentations, especially industrial ones, drives selection based on acidity tolerance gradually reducing the population diversity over the course of fermentation through directional selection.

## 2 Overview of vinegar production

Vinegar, in all its varieties, is primarily a solution of acetic acid in water accompanied by various other compounds depending on the raw material or additives. While acetic acid can be produced by industrial processes, all products termed ‘vinegar’ in most countries must be the result of biological fermentation. The fermentation organisms are acetic acid bacteria (AAB), a gram-negative rod shaped bacteria that belong to many different genuses and species in the family Acetobacteraceae. These bacteria are obligate aerobes and require oxygen to ferment acetic acid from ethanol. The basic equation is

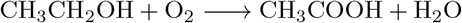

The reaction is actually two parts where one oxygen atom is combined with ethanol to make acetaldehyde and then subsequently another oxygen atom transforms acetaldehyde into acetic acid. The reactions are catalyzed by variations of the enzymes alcohol dehydrogenase and acetaldehyde dehydrogenase respectively. The reaction is also exothermic and requires heat to be removed from the process for large vinegar generators using heat exchangers or cooling towers.

One of the defining characteristics of many vinegar fermentations is the construction of a biofilm made primarily of cellulose that promotes bacterial colony formation and oxygen exchange with the atmosphere. It is most commonly known as the ‘mother of vinegar’. The strength of vinegar acidity is measured by titratable acidity which is defined as 1% acidity for every 1 gram of acetic acid per 100 mL of water. Most countries have a minimum legal acidity for vinegar of 4% while most store bought vinegars are usually 5% with the exceptions of wine vinegars or balsamic vinegars at 6-7%. While pH is also useful, it is not consistent across vinegars for the same acidities and is not used as a standard.

Even though vinegar has been made for centuries by many ‘slow’ methods that combine live vinegar mothers with alcohol such as wine or cider, these methods typically are laborious and take 2-3 months to complete. Starting in the early 19th century, industrial production methods were created which allowed for both more rapid fermentation but also higher acidities. The first method known as the ‘quick process’, invented by Karl Sebastian Schüzenbach in the Kingdom of Baden (now Germany’s Baden-Württemberg state), used layers of wood shavings inoculated with vinegar bacteria to increase the surface area for fermentation [11]. Combined with repetitive sparging and air flow this reduced production time to about 2 weeks. After World War II, Austrian engineers Otto Hromatka and Heinrich Ebner [12] adopted the submerged fermentation process recently developed for the mass production of penicillin to make vinegar. This new process of submerged fermentation is now the dominant technology for industrial vinegar fermentation globally.

Submerged fermentation functions by simultaneously injecting air and agitating a mixed mash of raw vinegar and alcohol with an impeller at the bottom of a tank. The air is dissolved and is used by the bacteria for rapid fermentation that can complete in as little as twenty-four hours. The final acidity can be high, up to 15% or more, in contrast to the 10% maximum for the old quick process and 5-7% for traditional slow methods. This is then diluted with water to retail strength.

Bacterial population dynamics in vinegar production typically have three main stages. First, a lag stage where the bacterial population selects species and strains that will survive in the new environment and begin fermentation. Second, an exponential phase where bacteria rapidly multiply and the fermentation rate accelerates. Third, is the carrying capacity stage where the population density stabilizes at the carrying capacity of the generator. This population then ferments vinegar at a nearly constant rate throughout the cycle.

### 2.1 Acetic acid bacteria and acid tolerance

AAB are nearly unique amongst all prokaryotes in their ability to tolerate and thrive in the high acidity environment of acetic acid. Both due to its acidity as well as unique chemical properties, even low levels of acetic acid (less than 0.5%) are lethal to most benign and pathogenic bacteria [13]. However, within the AAB types and strains within species, there are differential levels of tolerance to acidity that affects the population diversity of the bacteria throughout the fermentation cycle.

Acidity tolerance in AAB has many aspects including enzyme activity, mechanisms to pump acetic acid out of the intracellular environment and membrane proteins. Amongst the two most prominent enzymes facilitating the acetic acid resistance of AAB are citrate synthase, a prominent enzyme in the tri-carboxylic acid cycle of metabolism, and pyrroloquinoline quinone-dependent alcohol dehydrogenase (PQQ-ADH) [14–19]. The deletion of these enzymes is known to reduce or eliminate acetic acid resistance and expression data shows their genes are more highly expressed in high acid resistant strains.

However, there is differential acidity resistance amongst not just species but within species. The acetic acid environment that AAB create during fermentation causes selection pressures that drive the sweep of certain alleles through a population which allow higher acid production is a clear example of niche construction. This niche construction will be discussed and modeled to show how this effects the genetic diversity as well as average fitness over the course of the fermentation.

## 3 Niche construction in vinegar production

The niche construction in AAB during vinegar fermentation follows closely within the model outlined in equation 1. The environment here will be modeled as the acidity of the fermentation while the organism will be modeled as a population of organisms with alleles that convey varying levels of acetic acid tolerance to their organism. As the fermentation proceeds, the population evolves to consist only of those organisms that can tolerate high acidity which continue to raise the acidity, changing their environment.

### 3.1 Evidence for niche construction in AAB

The evidence for selection of AAB at higher levels of acidity has been well-founded in a variety of studies though it is often measured in terms of ecological succession amongst different species.

There are many genuses of acetic acid bacteria but two are most prominent in vinegar production. The genus Acetobacter contains bacteria that are dominant in the traditional modes of fermentation, most commonly *A. pasteurianus* and *A. orleanensis*. They typically have an acidity tolerance of 5-8%. The genus Komagataeibacter is more common in the industrial submerged fermentations of distilled ethanol (for distilled white vinegar), red and white wine, and cider vinegar. It can have an acid tolerance of 15-20% and is the workhorse of large-scale vinegar production.

### 3.2 Evidence for ecological succession and niche construction

There are many articles describing ecological succession throughout vinegar fermentation [20, 21]. Universally, they find a wide diversity of AAB in both genus and species in the initial stages at low acidity that gives way to increasingly lower diversity and a higher frequency of high acid tolerance strains as fermentation continues. Typically the transition is from a diversity of Acetobacter and Komagataeibacter to primarily Komagataeibacter. This demonstrates the evolution of the ecology with higher levels of acidity.

The evidence for evolution within species is more indirect. A common problem with bacteria from high acid fermentations is the difficulty of culturing them to allow analysis. Often the only bacteria that can be cultured show a lower acidity tolerance, based on acidity of culture media, than the vinegar they were sampled from implying high acidity tolerance AAB are different from those cultured. There is a plethora of evidence that high acidity tolerant AAB are not resilient to environmental conditions outside of the fermentation generator environment they have created. Submerged fermentation systems are very fragile and a loss of power for even only a minute can often destroy the culture. This is why such systems necessitate diesel generator or battery backups. Complex culture media with multiple layers are often needed to grow AAB from high acidity cultures and even these are not always successful [22].

However, despite the difficulty of culturing these AAB and subjecting them to genetic typing, there are several key pieces of evidence that point to low diversity within species in high acidity fermentations. First, in high acidity distilled ethanol and wine vinegar fermentations, those organisms that can be cultured are often of only one species [23]. In addition, analysis of plasmid sizes from samples of high acid fermentations indicate an increasingly smaller variety of plasmids at high acid fermentations that are replicated when a high acid fermentation is used to seed another generator and produce high acid vinegar [24, 25]. Finally, there is the well-known vulnerability of submerged fermentation systems to attack by bacteriophages.

Phage attacks are well-known in the vinegar industry as well as other bacterial fermentation food industries such as cheese manufacturing that use lactic acid bacteria to make cheese from milk [26–31]. Phage attacks are unheard of in traditional vinegar fermentation and usually less debilitating in the older quick process packed generators that produce lower acidities and have higher bacterial culture diversity [27]. Phage attacks on quick process systems typically slow production while they destroy the culture in submerged fermentation, sometimes within a few hours of introduction [27]. This is similar when monocultures of lactic acid bacteria are used to make cheese [31]. This vulnerability implies a very narrow number of highly related bacteria are present in high acid fermentations and the lack of genetic diversity leads to vulnerability to pathogens. Therefore, this implies that the environmental conditions the AAB create themselves lead to directional selection for species with higher acidity tolerance, even as this reduces the species and strain diversity.

In bacteria, since they are haploid organisms with a single chromosome plus plasmids, the effects of selection are typically represented as ‘periodic selection’ [32]. Periodic selection relates to the fact that the haploid nature of bacteria are such that a selected variant will allow the entire chromosome to essentially hitchhike as it is passed to future generations so long-range linkage disequilibrium and the clonal nature of bacteria after a selective sweep is common. Recombination does exist in bacteria, though it’s frequency varies by species [33], and is measured by the index of association, a multi-allelic linkage disequilibrium parameter [34]. However, the effects of periodic selection is such that bacteria that differ in multiple, putatively neutral, alleles can result from a selection event giving both a unique but also clonal population.

## 4 Niche construction model for vinegar fermentation

The niche construction for AAB in industrial fermentation is a positive feedback loop where acetic acid production by bacteria increases the total acidity, selecting out bacteria that cannot survive or produce well at lower acidities, and allowing an increasingly large proportion of the population to be occupied by those AAB that can both produce acetic acid at higher acidities and survive such environments. In this paper we will consider a uniform species of acetic acid bacteria that has multiple strains that differ by an allele variant that endows them with tolerance for acetic acid up to a certain acidity. For the purposes of this model we do not consider mutation or bacterial recombination that shares genetic material between strains and species. Plasmids are also not included since plasmids and their copy numbers have not been shown to be a key factor in acetic acid resistance unlike their crucial role in antibiotic resistance.

The model will assume the vinegar generator has ended the exponential phase and the bacterial population density is at carrying capacity and fermenting at its maximum rate. The population and its relative proportion by strain will fluctuate by modeling the dynamics of the bacteria reproduction using a branching process under the effects of both carrying capacity restraints and selection at varying levels of acidity.

### 4.1 Branching process reproduction model

The bacterial population dynamics will be generalized by a Bienayme-Galton-Watson branching process [35] that allows for three probabilities for an given bacterium: *p*_0_, the probability that the bacteria dies off and has no descendants, *p*_1_ the probability that the bacteria reproduces by binary division but only one organism survives to reproduce again and *p*_2_ the probability that the bacterium reproduces and both organisms survive to reproduce. The expected number of progeny per generation per bacterium, known as the reproduction number *m*, is given by

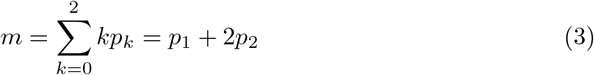

As is well-known in the theory of branching processes, if *m <* 1 the population tends to extinction with a probability of 1 over time. Otherwise, the population can grow exponentially to ‘infinity’ with a finite probability where *m* ≥ 1. Amongst a population of different strains of bacteria, the strain reproduction number is also a function of the relative fitness, *w*. The reproduction number, however, is not exactly equal to the relative fitness since it will consist of two components: a relative fitness component based on acidity and a general component, common across all strains, based on the population size in relation to carrying capacity.

The value of *m* can be related to both the carrying capacity and the tolerance for acidity. Assume that at carrying capacity and in an environment of tolerable acidity, *m* = 1. The two effects can be modeled as a product of the factor affecting overall population density and the fitness effects of acidity *A*, manifested through selection. The effect of carrying capacity, *K*, versus population density, *N*, will use the discrete time modification of the logistic equation to give

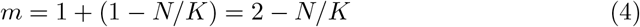

Likewise, the effects of acidity on the reproduction number can be given by *m*(1 − *s*(*A*)) where *s*(*A*) is the selection coefficient for a given level of acidity and *w*(*A*) = 1 −*s*(*A*) is the relative fitness of a strain at acidity *A*. This gives a new expression for *m*.

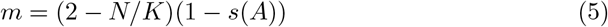

#### 4.1.1 Probability modeling with maximum entropy

Understanding the branching process and its expected value allows us to understand the general population growth or decline for a strain but does not directly gives us the probabilities *p*_0_, *p*_1_, and *p*_2_ needed to simulate the population. Here we will derive values for these probabilities assuming the probability distribution is the maximum entropy distribution given the constraints

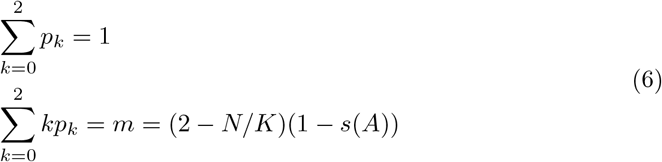

The entropy of the branching process probabilities is given by

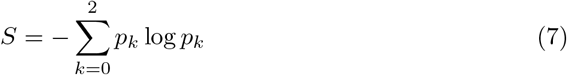

The derivation of these probabilities can be easily done using the Lagrange multiplier method. For a Langrangian function *L* and Lagrange multipliers *λ* and *μ* the Lagrangian is given by

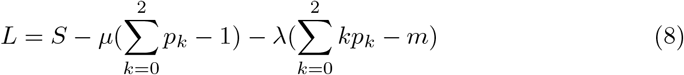

To find the maximum entropy probabilities we differentiate *L* with respect to *p*_*k*_ and equate it to zero giving

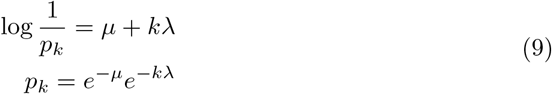

Since Σ *p*_*k*_ = 1 we can determine *μ* by

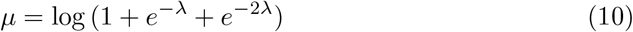

and *λ* can be determined by find the roots of the equation

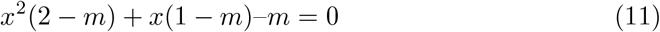

In the equation above *x* = *e*^−*λ*^ allows one to calculate *λ* once *x* is known. With both variables calculated then the branching process probabilities can derived.

Therefore, the probabilities of zero, one, or two descendants for a given group of bacteria are calculated based on the reproduction number. If population is near carrying capacity and the acidity is past tolerance, the reproduction number declines and the probability of *p*_0_ increases relative to the other probabilities.

### 4.2 The acidity relative fitness function

As stated previously, relative fitness, *w*(*A*) = 1 − *s*(*A*), is based on the acidity tolerance of a bacteria strain. The expression for *s*(*A*) is not elaborated in the literature but it is clear that past the acidity tolerance the bacteria begin to die off. Therefore, we will look at a fitness function where *s*(*A*) has different values below and above *A*^*^, the limit of acidity tolerance.

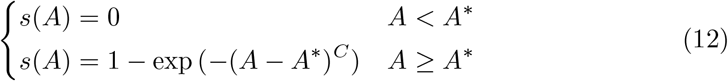

The variable *C* is an exponent that can be varied to increase or decrease the rate of selection against a strain once the acidity tolerance is reached. For this paper a value of *C* = 1*/*2 was selected.

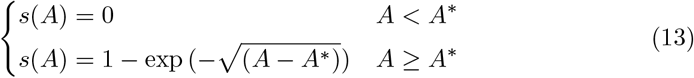

### 4.3 The acidity production function

The rate of acidity production by fermentation per population density, Δ*A* is given by another expression where the rate of acidity production is assumed to be the same for all strains relative to their acidity tolerance. Above the maximum acidity tolerance, the bacteria are too stressed to produce acetic acid and do not do so. Below the acidity tolerance, the acidity production depends on a constant, *α* which represents the base acidity production. As the acidity tolerance is approached, there is a slowdown in production that becomes more acute the closer the acidity is to the tolerance.

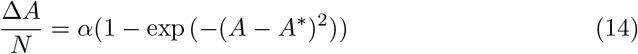

This function is not based on any explicitly biochemical framework or pathway since the relationship between acetic acid production and strain acidity tolerance is not yet clarified in the literature, however, it represents the behavior seen in actual fermentation where acidity production slows and then halts as the acidity tolerance for a strain is reached.

## 5 Niche construction coupled difference equations

Understanding how the rise in acidity is both driven by the bacteria as well as how it exerts selection and evolution of allele frequencies in the population we can derive the coupled difference equations following equation 2 where each strain is indexed by *i, E* = *A* and *O* = *f*_*i*_ where *f*_*i*_ = *N*_*i*_*/N* is the allele (strain) frequency.

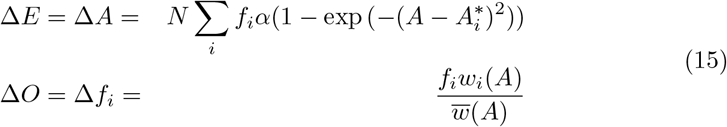

The expression for Δ*f*_*i*_ is the standard equation for the change of allele frequencies in haploid populations undergoing selection.

## 6 Simulation of vinegar fermentation niche construction

The expressions in Equation 15 are the governing equations involved the evolution of the niche construction, however, they are difficult to solve analytically due to the summation in the term for the acidity (environment). Therefore we will simulate the system to demonstrate the effects of niche construction.

In the simulation, a population of bacteria in an industrial fermenter, assumed to be at the steady state carry capacity population phase, will be simulated over time to demonstrate the effects of self-produced acidity on the bacterial genetic diversity. The original bacterial population *N*, will be established and divided across ten strains, originally of equal frequency *f* = 0.1. Each strain will have a different acidity tolerance, spaced equally over the range of starting acidity and the maximum acidity. Throughout the simulation the bacteria will produce acetic acid per equation 14 and as the acidity increases, strains will be selected against and become extinct.

The genetic diversity of the population will be measured using the effective number of species metric proposed by Jost [36] where the effective number of strains *D* is given by

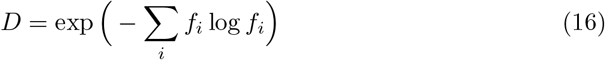

At each time step, total acidity, relative fitness, population size, and the effective number of strains will be measured.

The acidity of the vinegar will start at 4% and increase to a maximum of 15%. This is in line with production practices where new alcohol is combined with finished vinegar to have a starting acidity around 4% as well as enough live bacteria to rapidly reach the carrying capacity phase and begin steady fermentation. The value of *α* for acidity production is calibrated so that the total production will complete in about 24 hours given the population carrying capacity as is common in submerged fermentation. The value of the carrying capacity *K* and starting population *N* is 1,000,000. This is not an exact number of the amount of bacteria but rather a population density where each unit can contain thousands or millions of bacteria. Based on this value of *N*, a value of *α* is chosen as *α* = 4.6 *×* 10^−7^/(g/100mL)/bacterial unit/hour.

Figure 2 shows the results of the simulation. It is clear that while the population stays relatively high, the strain diversity progressively decreases with acidity at points where the acidity tolerance of each strain is exceeded. In addition are the sharp, but temporary dips in relative fitness as the acidity overtakes strains. Of key note is that as the acidity increases and the strain diversity declines, falls in relative fitness and bacterial population become more accentuated since the remaining strains are increasingly larger proportions of the overall population.

**Fig. 1.**
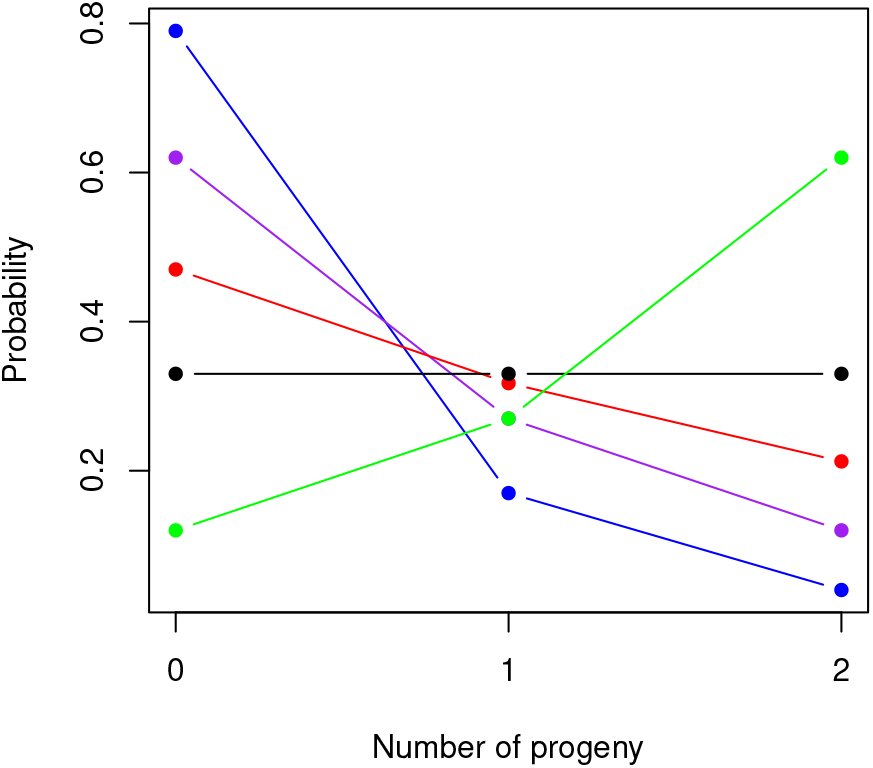
Maximum entropy probabilities for the Bienayme-Galton-Watson branching process governing bacteria reproduction for various values of *m*. Green is *m* = 1.5, black is *m* = 1, red is *m* = 0.75, purple is *m* = 0.5, blue is *m* = 0.25.

**Fig. 2.**
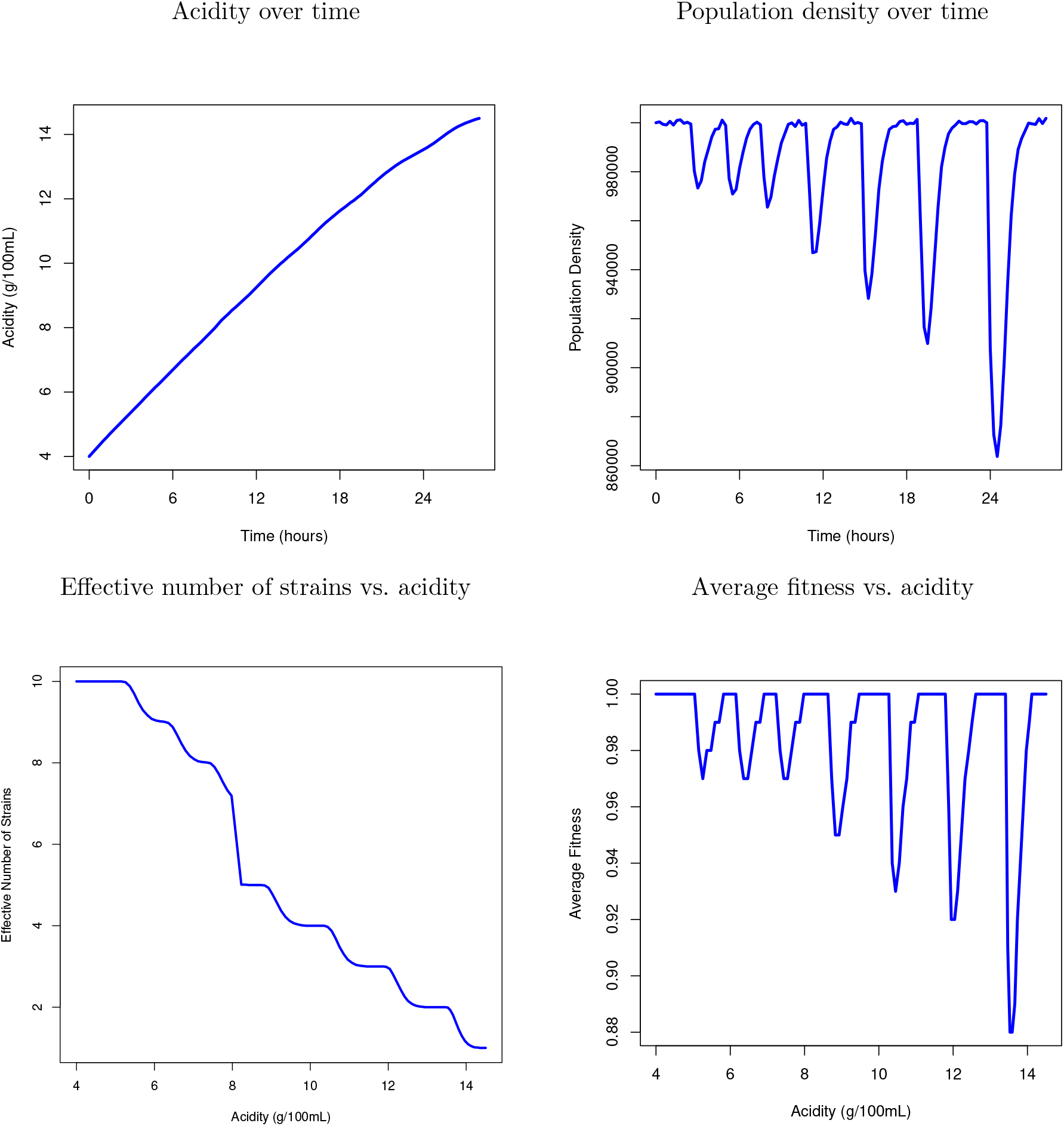
Simulation plots of niche construction by acetic acid bacteria. The top figures are the acidity over time (hours) as well as the population density over time. The bottom figures are the effective number of alleles (strains) at various levels of acidity and the average fitness at various levels of acidity.

## 7 Discussion

Industrial vinegar fermentation has strong evidence that it is a clear cut and readily quantifiable example of niche construction. Acetic acid production by the bacteria alter the environmental acidity and drive evolution of other loci in the population, particularly those loci on genes involved with acid tolerance. Due to the periodic selection nature of bacteria, many more loci are likely selected as well due to the chromosome-wide linkage disequilibrium. This could mean that niche construction in prokaryotes is much more consequential to evolutionary change since the effects at loci are not isolated by regular and frequent recombination. The ultimate impact would be that a few genes, likely responsible for secondary metabolites or fermentation, drive environmental change that allows wider selection for entire strains.

What is interesting about this niche construction is that its extreme manifestation is only possible in industrial fermentations. In nature, acetic acid bacteria are always present, for example feeding on the alcohol made by yeasts in rotting fruit, but the ultimate acidity is relatively low. There is speculation that the current industrially useful species possibly evolved from other acetic acid bacteria when selection pressures from the industrial production environment began in the 19th and 20th centuries, but this is still unproven. It is clear, however, that within production runs, evolution mediated by niche construction is not only key but of primary importance to an industry estimated to be at least $500 million per year in the United States alone and many times that worldwide. Without niche construction, large-scale and inexpensive production of vinegar may not be possible, so it is far from a theoretical concern.

Niche construction and selection of productive strains comes with risks, however, as outlined before. Research indicates increased industrial efficiency in many fermentation industries, dairy and vinegar to name two, are making bacteriophage attacks on production more common and possibly economically damaging though figures are not available. There has been speculation of similar phage issues in pharmaceutical industries that use recombinant bacteria in production but not much has been published, possibly due to industry secrecy or the higher requirements for sterility in pharmaceutical good manufacturing practices.

This also highlights, however, that niche construction is probably mediated in the wild by factors that don’t allow the extreme narrowing of diversity seen in controlled settings. The lower levels of acidity allow acetic acid bacteria to niche construct and eliminate other competing bacteria but not reduce species diversity to low levels. One could speculate many organisms have environmental or intrinsic factors that limit how narrowly a niche can be tailored. It is possible that in many niche construction scenarios, there ceases to be a significant relative fitness difference caused by the niche amongst organisms within the niche. Also, aspects of the niche itself could inhibit future niche construction. This allows the niche to have a development capacity that retains advantages for the resident organisms without damaging genetic diversity. A simplistic, though effective, metaphor is the limits from the Verhulst equation for logistic growth in contrast to the infinite exponential growth curve.

This would be interesting in many practical ways. In beneficial settings, like fermentation, raising or removing the limits of niche construction can lead to productivity, despite its risks. In other contexts, making the niche construction limits more stringent could inhibit the actions of pests or bacteria that secrete biofilms to improve their environment. This was directly addressed in [8] where antibiotic overuse in humans is modeled as a phenotype in a niche construction model. In these cases, while the organisms are not being addressed directly, their niche construction abilities can be modified to encourage certain outcomes. This would elevate niche construction into a practical tool that would emphasize the importance and usefulness of the paradigm.

## 8 Declarations

### Funding

The author declares that no funds, grants, or other support were received during the preparation of this manuscript.

### Competing Interests

The author has no relevant financial or non-financial interests to disclose.

### Data Availability

Source code is available from the author at request.

## Notes

### Competing Interest Statement

The authors have declared no competing interest.

